# Engineering GliaTrap: a biodegradable non-swelling hydrogel with tuned release of CXCL12 to attract migrating glioblastoma cells

**DOI:** 10.1101/2023.04.12.536581

**Authors:** Yusuke Suita, Saradha Miriyala, Deniz Merih-Toruner, Weizhou Yue, Lingxiao Xie, Blessing Akobundu, Nathan Pertsch, Andras Fiser, Eduardo Fajardo, Jie Shen, Nikos Tapinos

## Abstract

Glioblastoma is the most aggressive type of brain cancer with an average overall survival of 15-21 months after first diagnosis. The relapse is mainly caused by migrating glioblastoma cells that diffuse away from the tumor mass into the brain parenchyma and retain cancer stem cell (GSC) properties. Current therapeutic options are ineffective and inevitably result in relapse, indicating a high unmet medical need for innovative therapies in the treatment of invasive glioblastoma. To address this challenge, we propose a new therapeutic modality: GliaTrap, a biodegradable non-swelling, injectable hydrogel with sustained release of a chemoattractant for GSCs that lures and traps the migrating cells back to the tumor resection cavity. We developed a biodegradable and injectable hyaluronan/collagen II-based (HA/Col) hydrogel that does not swell in vivo. The hydrogel is embedded with CXCL12 loaded liposomes and is tuned for sustained release of CXCL12. The safety profile of liposome-embedded HA/Col hydrogel was determined in-vivo after stereotactic implantation in the mouse brain. The efficacy of GliaTrap to attract GSCs was determined ex vivo using a 3D tumor spheroid model and in-vivo using 3D light-sheet microscopy in orthotopic human glioblastoma xenografts. Our findings suggest that GliaTrap could represent a safe and efficacious new therapeutic approach for glioblastoma and potentially serve as a drug delivery platform to locally deliver tumor-killing agents.

**One Sentence Summary:** GliaTrap is a biodegradable non-swelling hydrogel with tuned release of a chemoattractant to attract invading glioma cells and serve as delivery platform for local therapeutics.

## INTRODUCTION

Glioblastoma (GBM) is the most malignant and aggressive grade IV primary brain tumor with an average overall survival of 16∼21 months after first diagnosis. Standard treatment of glioblastoma includes surgical resection followed by radiation in the vicinity of the resection cavity and administration of temozolomide (*1*). Even with this therapeutic approach, tumor recurrence is inevitable (*2*) primarily due to the presence of glioma stem cells (GSCs), which are characterized by high migratory potential (*3–5*), resistance to chemotherapy and radiation and the ability to form recurrent tumors (*6*). Eradication of the migrating GSCs remains an unmet medical need that leads to the high recurrence rate of glioblastoma.

To overcome this challenge, we engineered GliaTrap as a solution to sequester the migrating GSCs and potentially eliminate the cancer cells locally. GliaTrap is a biodegradable non-swelling hydrogel implanted into the tumor resection cavity after surgery and using a chemoattractant concentration gradient maintained by the hydrogel to lure residual GSCs to the vicinity of the resection cavity. To achieve the characteristics of GliaTrap, we used biopolymers that are abundant in the brain: collagen II and hyaluronic acid (HA). It has been shown that collagen-based in situ hydrogels can quickly form a gel at 37°C and were compatible with cells (*7, 8*). On the other hand, HA can help maintain cell phenotype (*9*).

The C-X-C motif chemokine ligand-12/C-X-C motif chemokine receptor-4 (CXCL12/CXCR4) signaling axis is involved in central nervous system (CNS) development (*10*), while in glioblastoma, it has been shown that it is important for invasion (*11*), GSC migration and therapeutic resistance (*12*). Due to its pro-migration properties, we decided to use CXCL12 as a chemoattractant for the GliaTrap proof-of-principle study. We developed our hyaluronic acid/collagen II-based (HA/Col II) hydrogel for the release of CXCL12, and characterized it to have non-swelling and injectable properties. To achieve sustained release of CXCL12 in the brain, we encapsulated CXCL12 into nano-liposomes and showed continuous CXCL12 release in an artificial cerebrospinal fluid (CSF) over several weeks. We showed that CXCL12 or the hydrogel alone do not induce any inflammatory effects in the brain in vivo hence they could be safely applied in an oncology setting. Using an ex vivo 3D engineered biomimetic environment we demonstrated that GliaTrap significantly attracts GSCs that are migrating away from a patient derived GSC tumor sphere. Finally, using X-CLARITY and 3D light-sheet microscopy we confirmed that GliaTrap significantly attracts patient derived GSCs from an in vivo human orthotopic xenograft glioblastoma model.

Cancer cell invasion away from the tumor bulk is one of the hallmarks of glioblastoma and significantly contributes to patient mortality. Here, we engineered GliaTrap that uses a chemoattractant gradient to revert the migratory stream of glioblastoma cells to confine them close to the tumor resection cavity. GliaTrap can be further engineered to provide a secondary release of a tumor killing agent to eliminate the cancer cells thus constituting a potential new therapeutic modality for human brain tumors and other solid malignancies.

## RESULTS

### CXCL12 induces GSC chemotaxis via CXCR4

CXCL12 was previously shown to induce the chemotaxis of glioblastoma cells (*13*). To confirm its functions as a chemoattractant for GSCs, we performed a chemotaxis assay in six patients derived GSCs using Incucyte live-cell imaging. Our results show that CXCL12 induces a significant increase in the migration index of GSCs compared to the non-treated group (Fig. 1A).

**Fig. 1.**
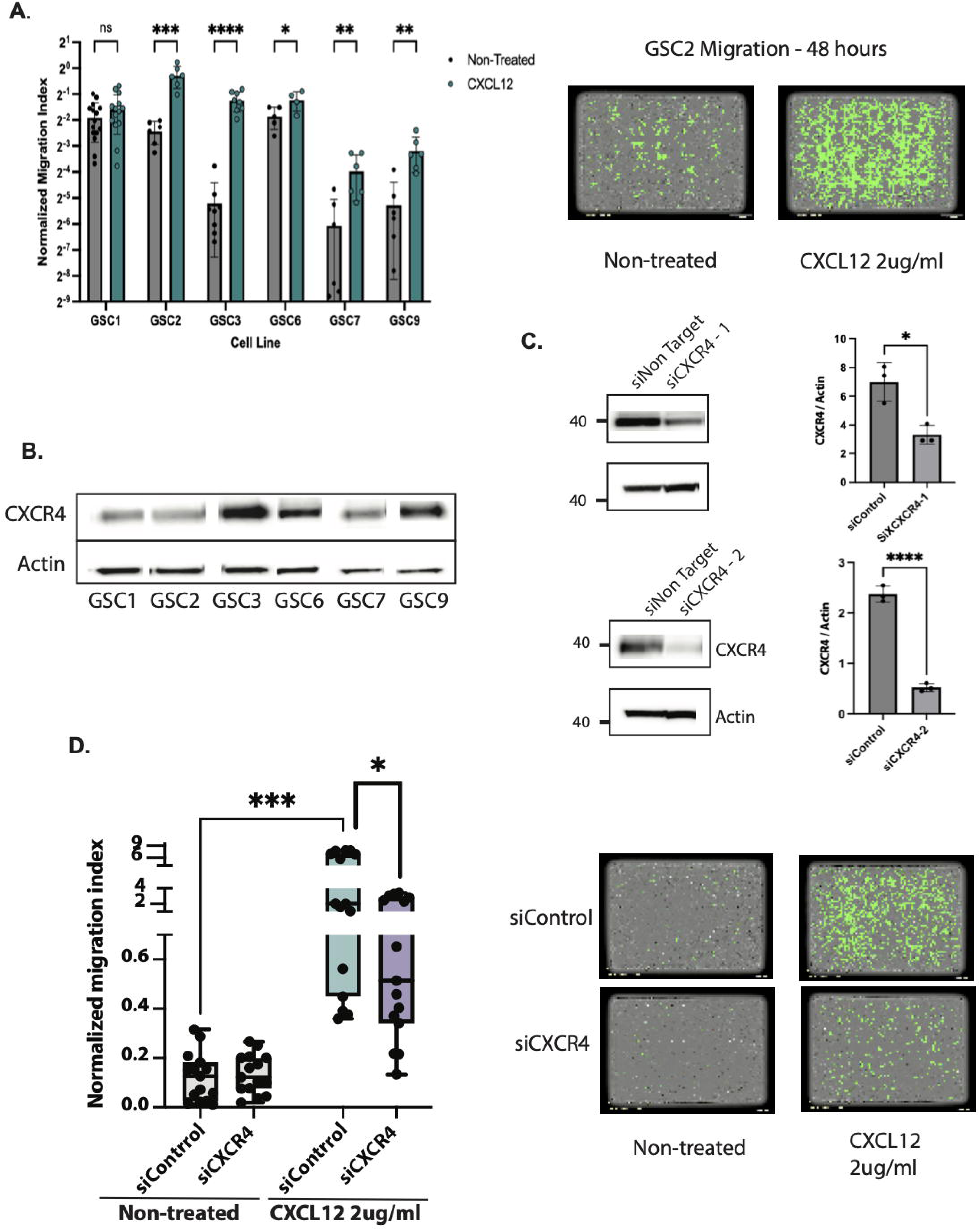
**CXCL12 induces GSC chemotaxis via CXCR4 receptor.** A) Chemotaxis assay of 6 patients derived GSCs treated with 2 ug/ml CXCL12. Images were taken at 48 hours using Incucyte Live cell imaging (Sartorius). Left panel shows normalized migration index ([GSC’s area in bottom well] / GSC’s area in top well)) at 48 hours compared to non-treated GSCs. CXCL12 induces significant increase in GSC migration (n=6-14 per GSC, *p<0.05, **p<0.001, ***p<0.00005, ****p<0.000001, Student’s t-test). Right panel shows representative images of migrating GSCs (objects with the green “mask”) in bottom well at 48 hours (left: non-treated, right: 2 ug/ml CXCL12). B) Expression of CXCR4 in 6 patients derived GSCs. C) Inhibition of CXCR4 expression with two independent siRNAs. Quantification of CXCR4 after siRNA inhibition shows that both siRNAs significantly inhibit CXCR4 expression (n=3, *p<0.05, ***p<0.0001, Student’s t-test). Actin was used as loading control. D) Chemotaxis assay of GSCs treated with 50 nM siCXCR4 for 3 days followed by incubation with 2 ug/ml CXCL12 for 2 days. Inhibition of CXCR4 induces significant inhibition of GSC chemotaxis in response to CXCL12 (n=12, ***p<0.001, *p<0.05, Student’s t-test).

CXCL12 has three known receptors CXCR4, CXCR7, and ACKR1 (*14, 15*). scRNA-seq data from the Brain Immune Atlas show that in glioblastoma the population of CXCR4+ cancer cells is about 60%, CXCR7+ cancer cells is about 2%, and ACKR1+ cancer cells are absent (*16*). Thus, we hypothesized that CXCR4 is the major receptor responsible for CXCL12-induced chemotaxis. To test this hypothesis, first we examined the expression of CXCR4 in six patients derived GSCs, which showed variable but universal CXCR4 expression (Fig. 1B). Then we inhibited the expression of CXCR4 with siRNAs (Fig. 1C) and measured the response of the CXCR4 knockdown (KD) cells on CXCL12 induced chemotaxis. This shows that CXCR4 KD induced a significant decrease in the CXCL12 induced migration index of GSCs (Fig. 1D). Finally, to exclude potential effects of CXCL12 on GSC proliferation, we treated GSCs with 2 ug/ml CXCL12 and performed an MTS assay. This shows that the growth rate of CXCL12-treated GSCs was equal to that of non-treated GSCs, indicating that CXCL12 does not induce GSC proliferation (Fig. S1A).

### Effects of CXCL12 on GSC transcript expression

To investigate the effect of CXCL12 on GSCs at the transcriptional level, we treated the GSCs with 2 ug/ml CXCL12 for two days and performed RNA-seq. Differential gene expression (DGE) analysis showed hundreds of differentially expressed genes, indicating that 2 ug/ml CXCL12 had a notable influence on transcripts in GSCs (Fig. S1B). To functionally classify the CXCL12 regulated transcripts, we performed gene set enrichment analysis (GSEA) on the DGE results using chemotaxis, EMT, and mesenchymal/proneural signatures. CXCL12-treated GSCs showed higher enrichment in chemotaxis and EMT signatures compared to non-treated GSCs (Fig. S1C). Mesenchymal signatures were enriched in CXCL12-treated GSCs while, proneural signatures were enriched in non-treated GSCs (Fig. S1D). To further investigate the effect of CXCL12 on transcriptional networks of GSCs, we performed weighted gene correlation network analysis (WGCNA). This analysis showed that CXCL12 transcripts correlate with CD248, SRGN, and ABI3BP genes, which are all responsible for cancer cell migration (Fig. S1E).

### Generation of a biomimetic 3D device to study effects of CXCL12 on GSC migration

To generate a biomimetic 3D device that could harbor human glioblastoma spheroids and could be used to study migration of GSCs, we utilized collagen I as an extracellular matrix component and polydimethylsiloxane (PDMS). The device was designed in a way that the chemoattractant is released from the left side of the device to form a concentration gradient, which could be detected by the glioblastoma cells migrating out from the GSC spheroid implanted in the center of the device (Fig. 2A). To ensure that our biomimetic device can be used to test the effect of CXCL12 concentration gradient on the directional migration of GSCs, we tuned the overall dimensions of the device and the shape and size of the posts that delineate the CXCL12 chamber from the center chamber that harbors the GSC spheroid. The final device has the following overall dimensions: width 16 mm, height 8 mm, and depth 3.175 mm. The width of the left side (CXCL12) and center (GSC) chambers were 3.682 mm and 7.364 mm, respectively. We selected circular over rectangular or hexagonal posts for the posts separating the CXCL12 chamber from the GSC chamber since this post shape supported the best linear gradient effect. Finally, we tested different numbers of posts and different distances between posts and determined that six posts with 0.653 mm distance between the posts provides the ideal environment for CXCL12 linear gradient formation. We used our optimized device with fluorescent dextran (10 kDa), which is a fluorescence molecule with a similar size to CXCL12, in the left chamber of the device and observed the release of dextran into the collagen I environment in the center chamber for 42 hours (Fig. 2B). This showed a linear gradient formation and continuous release of dextran into the center chamber suggesting that this architecture of the device is suitable for studying the effects of CXCL12 on GSC migration.

**Fig. 2.**
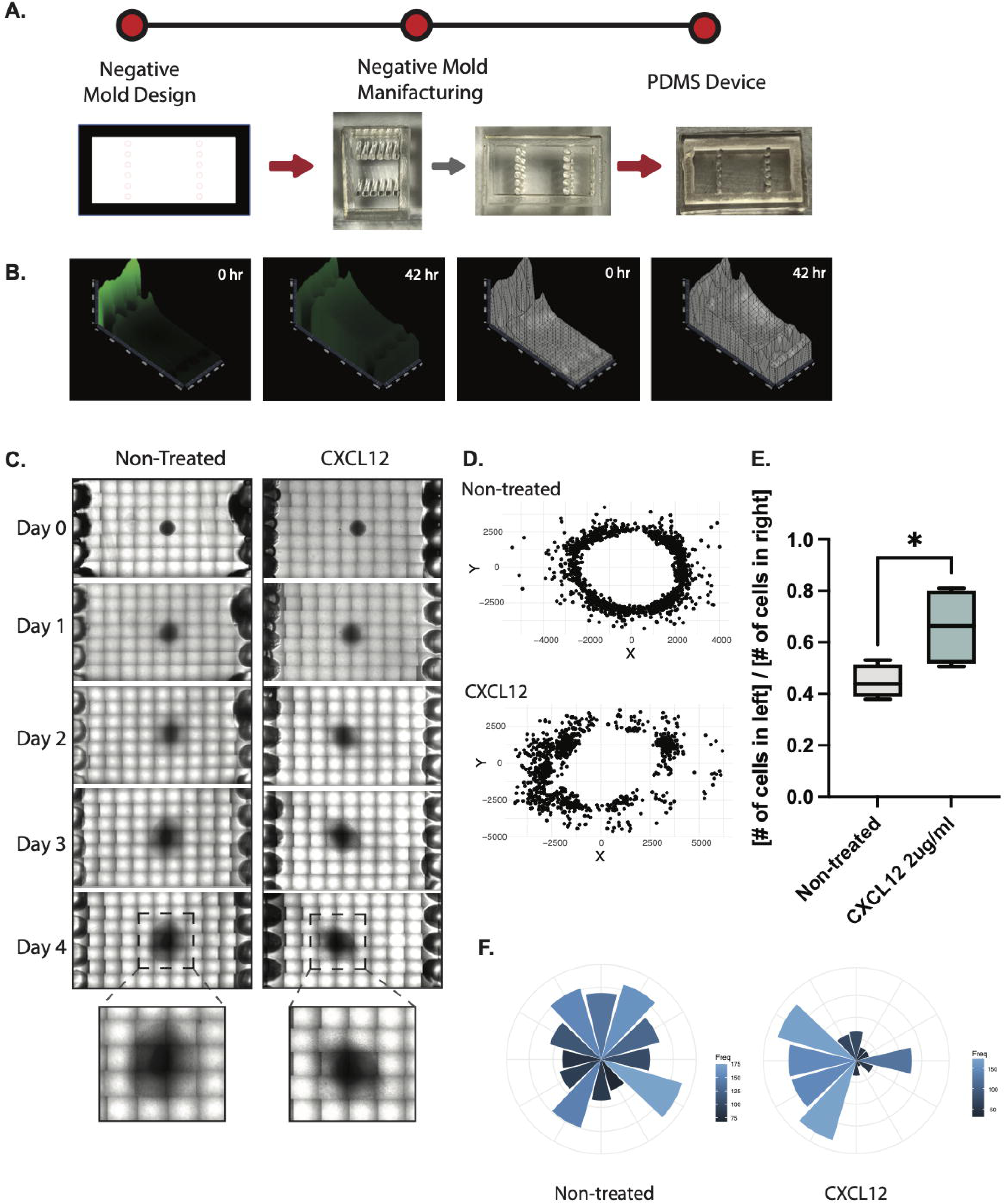
**CXCL12 induces GSC chemotaxis in a biomimetic device** A) Design of the negative mold for the biomimetic device and development of the PDMS device. B) Time-lapse images between T=0 and T=42 hours for dextran release from our hyaluronic acid/collagen II-based (HA/Col) hydrogel in the device using Zen 2.5D visualization software (Zeiss). C) Time-lapse images of GSC migration in the device in the presence or absence of GliaTrap+CXCL12. Images were acquired each day for 4 consecutive days. GSC spheroids were embedded in the middle chamber of the device in Collagen I. D) Coordinates of each migrating GSC in control or GliaTrap devices at day 4 were plotted using R. E) Quantification of the ratio of migrating GSCs between the left and right side of the middle line of each GSC spheroid. GliaTrap induces significant increase in the number of GSCs migrating towards the GliaTrap containing chamber of the device (*p<0.05, Student’s t-test). F) Coordinate single cell migration data were used to calculate the polar coordinates and degrees from the origin and visualized as a rose plot using the R package ggplot2.

### CXCL12 concentration gradient attracts GSCs migrating from a glioblastoma spheroid

Using our biomimetic 3D device, we tested the effect of CXCL12 on GSCs migrating out from a tumor spheroid. We transferred 2 ug/ml CXCL12 in the collagen I mixture in the left chamber of the device and a GSC tumor spheroid in the center chamber of the device and observed the GSC migration for four days using live cell microscopy (Fig. 2C). To quantify the effect of CXCL12 on GSC migration, we measured the number of GSCs that migrated towards the left and the right chambers after four days and calculated the ratio of cells towards the left versus the right. This shows that CXCL12 induces a significantly higher ratio of cells migrating towards the CXCL12 chamber (left) as compared to the negative control (non-treated) (Fig. 2D & E). Then we mapped the angles and coordinates of each individual cell migration in relation to the GSC tumor spheroid and quantified the effect of CXCL12 on the directionality of cellular migration. This quantification showed that CXCL12 induces a significant increase in directional migration towards the CXCL12 chamber (Fig. 2F) validating that a CXCL12 concentration gradient preferentially attracts GSCs migrating out from a glioblastoma tumor spheroid.

### Characteristics of the HA/Col II hydrogel containing liposome encapsulated CXCL12

To maintain protein stability and achieve sustained release of CXCL12 from the hydrogel, we encapsulated CXCL12 in liposomes and formulated a CXCL12 liposome embedded hydrogel. CXCL12 loaded liposomes were 217.1 ± 20.45 nm in size with a particle size distribution (PDI) of 0.353 ± 0.022 and neutrally charged with a zeta potential around 0 mV. The CXCL12 loading efficiency was 99.8%. The liposomes were mixed with the hydrogel materials prior to the gelation process. As shown Fig. 3A, the liposome-HA/Col II hydrogel mixture was in a liquid state at room temperature and gelled quickly at 37°C in approximately two minutes, through crosslinking between succinimidyl groups of PEG and amine groups of collagen II. The storage modulus of the liposome-HA/Col hydrogel composite was 234.41 ± 9.09 Pa, which is comparable to that of many soft tissues including the brain (∼140-620 Pa) (*17*), suggesting that the developed liposome/hydrogel composite may be well tolerated in the brain tissues. The swelling property of the HA/Col II hydrogel and liposome/hydrogel composite were also investigated. The swelling ratio of the HA/Col II hydrogel and liposome-HA/Col II hydrogel in the artificial cerebrospinal fluid at 37°C was 5.18% ± 0.46% and 4.39% ± 0.69%, respectively (Fig. 3B). This confirmed that our hydrogel system had low swelling in the aqueous environment. More importantly, the liposome-HA/Col II hydrogel demonstrated sustained release of CXCL12 over two weeks with a steady daily payload release (Fig. 3C).

**Fig. 3.**
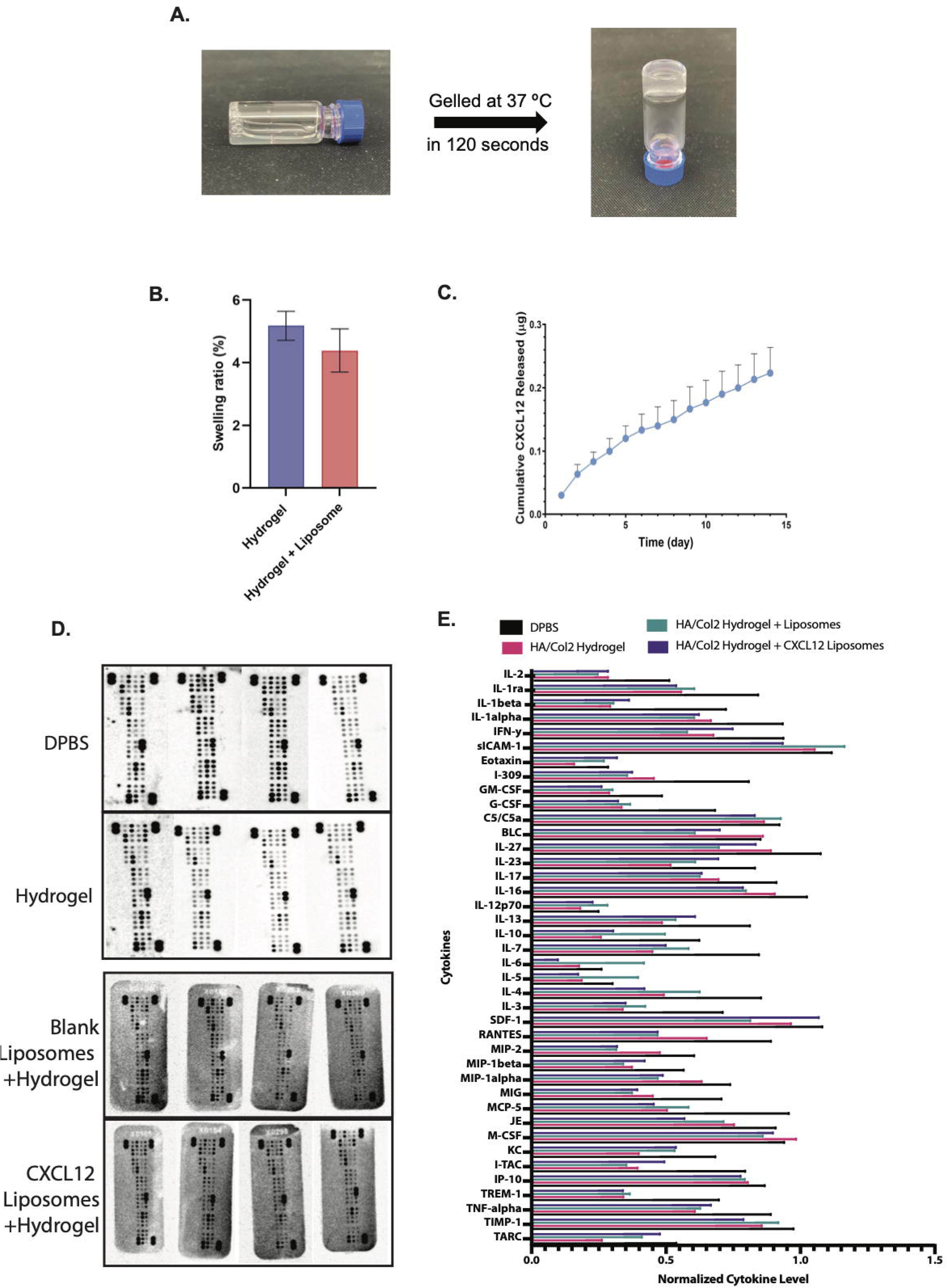
GliaTrap composed of hyaluronic acid/Col II hydrogel does not swell, provides sustained release of CXCL12, and does not cause inflammation in vivo. A) The gelation time of GliaTrap was determined using a tube inversion method at 37°C. B) The swelling ratio of GliaTrap was studied in an artificial cerebrospinal fluid. GliaTrap shows low (up to 5%) swelling. C) In vitro CXCL12 release kinetics from GliaTrap show sustained release of CXCL12 over 15 consecutive days. The cumulative amount of CXCL12 was measured by ELISA. D & E) Cytokine arrays using brain lysates from control mice (DPBS), mice with implanted HA/Col II hydrogel only, mice with implanted HA/Col II hydrogel + Blank liposomes and mice with implanted GliaTrap (HA/Col II hydrogel + CXCL12-loaded Liposomes). Quantification of the amount of each cytokine of each group was performed by calculating [signal intensity of a dot for each cytokine] / [signal intensity of reference dot] and comparison were performed with Anova. GliaTrap does not have any effect on pro-inflammatory cytokine levels in the brain.

### CXCL12 liposome-embedded HA/Col II hydrogel has no inflammatory side effects in vivo

To verify the gelation of HA/Col hydrogel at body temperature (37°C), we created an in-vitro environment that mimicks the stiffness and temperature of the mouse brain by pre-warming Matrigel to 37°C (Fig. S2A). Afterward, we loaded fluorescent dextran (10 kDa) into a liquid HA/Col II mixture and injected the dextran-loaded mixture into the pre-warmed Matrigel using the same injection method as that in the in-vivo experiment. We confirmed that dextran diffusion of the HA/Col II hydrogel group was much slower than that of the dextran in DPBS group (Fig. S2B), indicating that the HA/Col II hydrogel becomes a gel quickly at 37°C, the body temperature of the mouse.

CXCL12 has been shown to exhibit both pro- and anti-inflammatory properties (*18–20*). To examine whether the CXCL12 released from our hydrogel, induces inflammatory reaction in the brain, and to exclude the possibility that our hydrogel itself induces any inflammatory response, we performed stereotactic injections of the CXCL12 liposome embedded HA/Col II hydrogel, blank liposome-embedded HA-Col II hydrogel, HA/Col II hydrogel only and DPBS as a control in the brain of C57/B6 immune competent mice. One week later, we harvested the mouse brains (Fig. S2C) and performed mouse cytokine array analysis on the brain lysates. The result showed that the CXCL12 liposome-embedded HA/Col II hydrogel, the blank liposome-embedded HA/Col II hydrogel and the HA/Col II hydrogel did not significantly affect cytokines compared with the DPBS control group (Fig. 3D & E). This confirms the biocompatibility of the liposome-embedded HA/Col II hydrogel for in vivo applications and suggests that gradual release of CXCL12 from the liposome-embedded HA/Col II hydrogel does not elicit inflammatory responses in the brain and therefore, can be used as the hydrogel system for GliaTrap.

### GliaTrap disperses along blood vessels in the brain

Since GliaTrap is thermoactivated, injected as a liquid, and becomes a gel at body temperature (Fig. S2A), we examined if injection of GliaTrap in the brain parenchyma could form gel projections along preexisting structures in the brain like blood vessels. We loaded fluorescent dextran as shown above and injected the dextran loaded GliaTrap in the brain of Balb/C mice (n=3). One hour after the injection, mice were sacrificed and the brains were processed for tissue clearing using X-CLARITY a lipid-targeting tissue-clearing technique, which has been used to study cell movement in the mouse brain at a single-cell resolution level due to its tissue-clearing quality and relatively fast runtime (*21, 22*) (Fig. 4A). Following tissue clearing the brains were stained for smooth muscle actin to label blood vessels and imaged with 3D light sheet microscope (Olympus). This showed that GliaTrap forms several projections within the brain parenchyma following existing blood vessel formations (Fig. 4B, arrows) and even covers some blood vessels in the vicinity of the injections site (Fig. 4B, inset arrowheads).

**Fig. 4.**
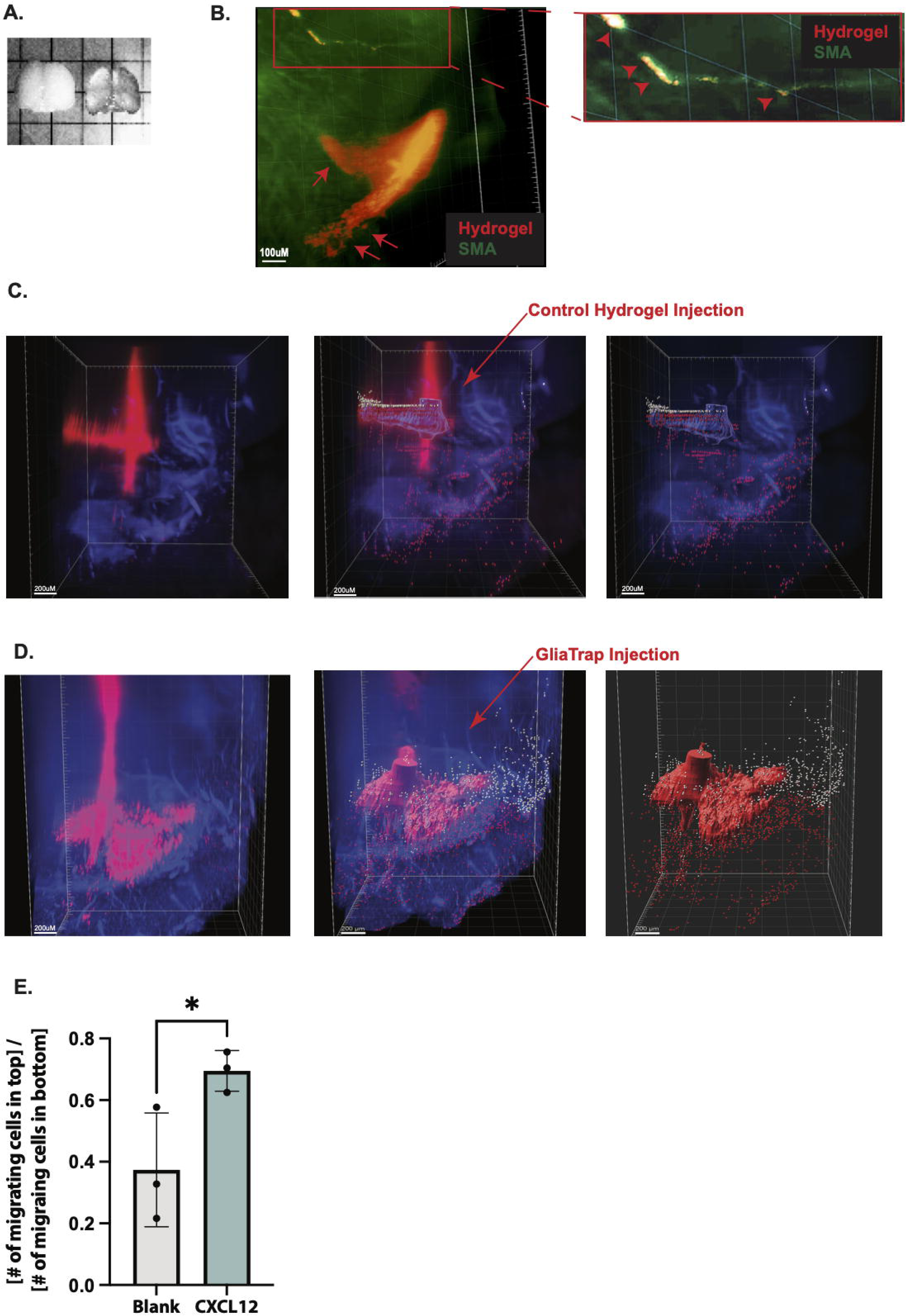
**GliaTrap disperses along blood vessels in the brain and attracts migrating GSCs in vivo** A) Representative image of a mouse brain processed with CLARITY, a tissue clearing technique (left: mouse brain with cerebellum area removed before CLARITY, right: the same mouse brain after CLARITY). B) Representative 3D-light sheet image of dextran-contained GliaTrap (pseudo-colored red) showing gel projections along pre-existing blood vessels stained with smooth muscle actin (arrows). Inset shows a magnified image of a brain blood vessel (green) covered with the GliaTrap hydrogel (arrows). C) Representative 3D light sheet images of the control group of mice (n=4). Human glioblastoma tumor (red) and single glioblastoma cells migrating mainly below the tumor mass are shown. Arrow depicts the area where the control (blank) hydrogel was injected under stereotactic guidance. D) In mice implanted with GliaTrap (n=4), single glioblastoma cells are migrating below (red) and above (white) the tumor mass. Arrow depicts the area where the GliaTrap was injected under stereotactic guidance. All images were processed using Imaris Imaging Software. E) Quantification of the ratio of migrating cells above the center line of the tumor mass to those below the center line shows that GliaTrap induces significant increase in the number of migrating glioma cells towards the GliaTrap injection site above the tumor mass, confirming the chemoattracting properties (n=6, *p<0.01, Student’s t-test).

### GliaTrap attracts migrating GSCs from glioblastoma tumors in vivo

To investigate the efficacy of GliaTrap (CXCL12 liposome-embedded HA/Col II hydrogel) in vivo, we injected ZsGreen-expressing human GSCs into eight Nu/J mice at coordinates of −2.0 mm AP and +1.5 mm ML relative to Bregma and -3.5 mm DV. These coordinates reflect the anatomic location of the putamen (sub-hippocampal region). One week later, we implanted GliaTrap into the brains of four mice and empty liposome-embedded HA/Col II hydrogel into the other four mice at 1.0 mm above the tumor injection site and monitored these mice for an additional week. This injection coordinate was selected because anatomically is in the cortical region above the hippocampus and glioblastoma cells are usually not migrating through the hippocampus (*23*). We posited that if human GSCs migrate through the hippocampal region, it will provide stronger evidence for the power of GliaTrap to attract migrating cancer cells. During the one week of monitoring after the implantation of GliaTrap, no motor deficit nor other abnormal behavior was observed, indicating that the GliaTrap did not induce any neurological damage either by hydrogel swelling or inflammatory reaction. A week after GliaTrap implantation, we harvested the brains and performed X-CLARITY. Following X-CLARITY, the processed brains were imaged using a 3D light-sheet microscope (Olympus). To measure the human GSCs migrating away from the tumor mass, we first defined the tumor mass area and the migrating GSCs based on the local maxima of signal intensity (Fig. 4B & C). To show preferential migration of human GSCs from the tumor mass towards GliaTrap, we measured the number of migrating GSCs above and below the tumor mass-defined region. We observed that GliaTrap induced a significant increase in the ratio of cells migrating at the area above the tumor where GliaTrap was implanted relative to the region below the tumor (Fig. 4C & D).

## DISCUSSION

In situ gelling hydrogels have attracted considerable interest in the past decade, owing to their advantages over pre-formed rigid polymeric implants (e.g., Gliadel^®^). These advantages include: 1) excellent injectability with minimal invasiveness; 2) ability to conform to the available space or reaction cavity at the injection site to provide maximal hydrogel/tissue interface allowing optimal payload access to the tumor tissues; and 3) tunable mechanical properties that are well tolerated by the surrounding brain tissues. The HA/Col II hydrogel is mainly composed of natural biopolymers collagen and HA and hence possesses superior biocompatibility compared to synthetic biodegradable polymers such as poly(lactic-co-glycolic acid) which degrades into monomers that acidify the local environment leading to potential local inflammatory reaction (*24*). It is noteworthy that our hydrogel system had low swelling in the aqueous environment which is ideal for maintaining good compatibility with surrounding tissues, particularly in the brain, as demonstrated in the present research using a mouse model.

The characteristic finger-like invasion pattern of glioblastoma cells around existing brain structures (myelinated fibers, blood vessels, axonal tracks) was first described by Scherer (*25*) in the 1930s. CXCL12 is highly expressed in these structures and contributes to the induction of migration of CXCR4-positive glioblastoma cells (*26–28*). The ability of CXCL12 to guide GBM cell migration has been supported by in vitro experiments (*11, 29*), however the molecular mechanisms involved are not completely defined (*30, 31*). In addition the role of CXCL12 in modulating the activity of CXCR4 and CXCR7 in GSCs (*32*) as well as the function of CXCL12 to mediate radio-resistance of GSCs in the subventricular zone (*33*) have been documented. These studies, although confirm the potent pro-migratory and chemoattractant role of CXCL12 for GSCs, they also suggest an adverse pro-tumorigenic role. Accordingly, to safely employ CXCL12 as the chemoattractant for GliaTrap requires controlled release both in terms of concentration and duration. Here, we showed that GliaTrap can achieve tuned release of CXCL12 over weeks and that in vivo application of GliaTrap releasing CXCL12 in the brain does not induce any pro-inflammatory reactivity from the microenvironment. To maximize the effectiveness of GliaTrap and to mimic Scherer structures in the brain by creating artificial finger-like projections of the hydrogel, we made GliaTrap thermo-reactive. GliaTrap is initially injected as liquid at room temperature; and can disperse in the interstitial space of the brain where it polymerizes to form hydrogel finger-like projections in reaction to the increased body temperature. Since the hydrogel projections of GliaTrap contain liposome-encapsulated CXCL12, they mimic Scherer structures to provide a more physiological environment to attract migrating glioma cells expressing CXCR4.

Our work proves that GliaTrap containing CXCL12 efficiently attracts a significant portion of migrating glioma cells in orthotopic in vivo models of human glioblastoma, however it does not address how these attracted glioma cells could be eliminated. Taking advantage of the liposomes embedded within GliaTrap, we are currently developing a dual release system where the initial release of CXCL12 is followed by a secondary release of therapeutics embedded in liposomes with slower release kinetics. Another option currently under investigation is the use of energy transfer through focused high frequency ultrasound and induction of thermal ablation of cancer cells in contact with GliaTrap projections. In conclusion, we envision GliaTrap as a platform technology that could attract and deliver various therapeutics (mRNA, antibodies, siRNAs, small molecules etc.) in the brain and other solid tumors.

## MATERIALS AND METHODS

### Primary glioblastoma stem cell isolation and culture

The IRB of Rhode Island Hospital has approved the collection of patient-derived glioblastoma multiforme tissue. All collections were performed with written informed consent from patients in completely de-identified manner and the studies were performed in accordance to recognized ethical guidelines (Belmont Report). Primary GSC spheres were cultured from human glioma samples as previously described (*34*). GSCs used in this study were authenticated by ATCC using short tandem repeat (STR) analysis. GSCs used were between passages 5 and 30 and cultured either as spheres or as attached on fibronectin-coated plates (10 ug/ml, Millipore Sigma) in a medium of 1X Neurobasal Medium, B27 serum-free supplement, minus Vitamin A, 100X Glutamax (Fisher Scientific), 1 mg/ml Heparin (STEMCELL Technologies), 20 ng/ml epidermal growth factor (Peprotech), 20 ng/ml basic-fibroblast growth factor (Peprotech). All GSC cultures are routinely tested for mycoplasma contamination using the MycoSensor qPCR assay (Agilent).

### Lentiviral transduction of glioblastoma stem cells

Lentivirus containing pHAGE-EF1a-ZsGreen-IRES-puromycin plasmid was concentrated by ultracentrifugation at 23,200 rpm for 90 minutes, then virus pellets were suspended in HBSS (Gibco). The virus titer was determined with Lenti-X™ p24 Rapid Titration ELISA Kit (Takara), according to the manufacturer’s instructions. GSCs (1 x 10^5^/12 well) were transduced in condition media by spin-infection (940xg) at 32°C for 2 hours. Puro-resistant and ZsGreen-positive GSCs were generated after seven days of puromycin selection, and the purity of stable GSCs was confirmed by flow cytometry analysis.

### In-vitro live-imaging chemotaxis assay

Inserts and reservoirs in an IncuCyte ClearView migration plate (Essen BioScience, 4582) were coated with a solution consisting of 10 ug/ml fibronectin (Sigma, FC0105MG) to 1% BSA (Fisher Scientific, NC0582624) in HBSS (Gibco, 14025076) at 37°C for 30 minutes. GSCs were resuspended at a concentration of 66,666 cells/ml in NBA complete media. After aspiration from inserts and reservoirs, a 60ul cell suspension containing 4,000 cells was seeded into the inserts and allowed to rest at room temperature for 15 minutes. Complete media or complete media containing 2 ug/ml of CXCL12 (Fisher Scientific, 350-NS-050) (200 ul) was added to the appropriate reservoirs. The IncuCyte ClearView cell migration plate was placed into the IncuCyte instrument SX1 (Essen BioScience) and scanning with a 10x objective lens for every 2 hours was scheduled over 120 hours according to the manufacturer’s instructions. Normalized migration indices were calculated by normalizing the area of cells in the reservoir at 48 hours to the initial area of cells in the insert.

### In-vitro proliferation assay

In-vitro proliferation assay was performed using MTS assay. GSC 4,000 cells were treated with CXCL12 2 ug/ml for five days in complete culture media (200 ul) in Nunc™ MicroWell™ 96-Well, Nunclon Delta-Treated, Flat-Bottom Microplate (ThermoFisher) pre-coated with fibronectin. Five days later, MTS reagent 20 ul (Abcam, ab197010) was added to each well, and the absorbance at 490 nm was measured every 30 minutes for 4 hours at 37°C. Absorbance at 490 nm was plotted using GraphPad Prism version 9.4.1. Cell morphology was observed every 2 hours during the five days using IncuCyte® live-cell imaging.

### siRNA transfection

GSCs were grown in a fibronectin-coated plate and transfected with 50 nM siRNA targeting CXCR4 (Dharmacon, L-005139-00-0005) or a non-targeting control siRNA (Dharmacon, D-001810-10-05) by using Opti-MEM (Gibco, 31985062) and the TransIT-X2^®^ Dynamic Delivery System (Mirus, 6000) in complete media without heparin (StemCell Tech, 07980) and Anti-Anti (Fisher Scientific, 15240062). The media was replaced with complete media without Anti-Anti 24 hours after transfection. The cells were incubated for 48 more hours before they were lysed or used in subsequent experiments.

### Western blot analysis

Cell lysates were prepared in 1x SDS lysis buffer containing 10 mM protease inhibitor cocktail (Millipore Sigma, P8340-1ML) and quantified using the Qubit Protein Assay Kit (Fisher Scientific, Q33211). Protein samples (20 ug) were used to perform electrophoresis in 1x Bolt™ MOPS SDS Running Buffer (Fisher Scientific, B0001) in Tris-buffered saline containing Tween-20 (TBST) (Fisher Scientific, 28360) in a Bolt 4-15% Bis-Tris Plus gel (Fisher Scientific, NW04120BOX) following the manufacturer’s instructions. The antibodies were incubated with primary antibody anti-CXCR4 (1:1000 in 5% BSA) (Abcam, ab181020) or β-actin (1:10000 in 5% milk in TBST) (Sigma, A1978) according to the manufacturer’s instructions. The membranes were then incubated with HRP-conjugated secondary antibodies (Cell Signaling Technology) at room temperature for 1 hour. The protein signals were detected using the chemiluminescent kit Radiance Q (Azure Biosystems AC2101). The western blot images were quantified by FIJI software (*35*) and normalized to the corresponding β-actin control.

### RNA sequencing and analysis

The 5x10^6^ attached GSCs were treated with recombinant CXCL12 2 ug/ml for two days. GSCs were lysed using the Trizol reagent (Invitrogen). RNA was isolated from this lysate using the RNeasy mini kit (Qiagen). Next-generation sequencing was performed on these RNAs. Sequence reads were aligned to the human hg38 genome assembly with hisat2 (*36*). Read counts were summarized with FeatureCounts using the refseq annotations found in the refGene.txt file (http://hgdownload.cse.ucsc.edu/goldenPath/hg38/database/refGene.txt.gz) (*36*). Protein-coding genes were selected, and differential gene expression analysis was performed on these protein-coding genes using DEBrowser, a R package to detect changes in gene expression (*36*). Eseq2, a R package, was used to perform differential gene expression analysis (*37*). Genes with fold change>1.5 and p-value<0.05 were considered for functional analysis.

### Gene Set Enrichment Analysis

Gene Set Enrichment Analysis (GSEA) was performed on the outcome of differential gene expression analysis using GSEA (Version 4.1.0) (*38*) to see if particular sets of genes are enriched, such as chemotaxis, metastasis, proliferation, DNA damage, mesenchymal GBM, and proneural GBM.

### Weighted Gene Correlation Network Analysis (WGCNA)

Weighted Gene Correlation Network Analysis (WGCNA) was performed using the WGCNA R package (Version 1.70-3) to identify gene and module networks that have similar gene expression patterns with CXCL12. Particularly, WGCNA was performed on publicly available GSC data (*39*). Genes from the modules with the highest correlation and p-value < 0.05 were picked to show the CXCL12-associated network. This network was visualized using Cytoscape (Version 3.8.2) (*40*).

### PDMS device design

The PDMS device was designed and manufactured for testing the CXCL12 concentration gradient by using a previously established method. The device’s dimensions are width of 16 mm, height of 8 mm, and depth of 3.175 mm. The device is designed with round edges and has six posts on each side with diameters of 0.635 mm, and the distance between posts is 0.653 mm.

### Chemoattractant kinetics imaging assay

Dextran (Thermo Fisher Scientific, D7170) 2.5 ug with collagen I mixture (100 ul) was loaded into the left chamber of the PDMS device. 0.1% BSA-containing DBPS in Cultrex rat Collagen I solution (R&D Systems, 3440-005-01) 170 ul was loaded in the center chamber, and 100 ul was loaded to the right chamber of the device. The dextran-containing collagen I mixture was polymerized at 37°C first to prevent the release of dextran into the center chamber. The collagen mixture was loaded into the center and right chambers and then polymerized at 37°C. After the polymerization, the device was imaged using Zeiss Apotome 2 with 10x magnification and tile mode. This procedure was done every 3 hours for three days. The release was visualized using Zeiss imaging software Zen’s 2.5D visualization mode.

### 3D model

The PDMS device is placed in a non-treated 6-well plate (USA Scientific, CC7672-7506) along with DPBS-soaked Kimwipe to avoid evaporation. One day before this GSC sphere invasion experiment, 100,000 GSCs in 200 ul were plated in a 96-well untreated round bottom plate (Corning, 351177). The plate was centrifuged at 110 rpm for 10 minutes at room temperature and was incubated at 37°C 6% CO_2_ for 24 hours to allow the formation of a GSC sphere.

CXCL12 (R&D Systems, 350-NS-050), dissolved in 0.1% BSA in DPBS, in Cultrex rat Collagen I was added to the left side of the device, and 0.1% BSA-containing DBPS in collagen I solution was added to the right side of the device. After collagen I polymerization at 37°C for 1 hour, a GSC sphere was placed in NaOH-stabilized Cultrex rat collagen I (R&D Systems, 3440-005-01). To prevent incomplete and premature gelation of the collagen, the 60 mm untreated 6-well plate, the Eppendorf tubes, and the pipette tips were pre-cooled. The plate was then imaged using the Zeiss Apotome 2 with 10x magnification and tile mode. This procedure was done every 24 hours for five days.

The number of migrating cells on both left and right sides of the sphere was quantified by FIJI. The final coordinate data was used to calculate the polar coordinates and degrees from the origin and visualized as a rose plot using the R package ggplot2 (ver. 3.3.6).

### Preparation and Characteristics of HA/Col II hydrogel

CXCL12-loaded liposomes were formed by mixing Lipofectamine 2000 (Thermo Fisher) and carrier-free human CXCL12 protein (R&D systems) at 1:1 molar ratio and the mixture was incubated at room temperature for 2 hours. The particle size, size distribution (polydispersity index, PDI), and surface charge of liposomes were characterized using Zetasizer Nano ZS90 (Malvern Instruments Ltd., Malvern, UK). The backscattering angel was 173° with a standard laser wavelength λ of 633 nm. The measurements were carried out in triplicate. The CXCL12 loading efficiency (LE) was determined using an ultra centrifugal filter (Amicon^®^-0.5, 30 kDa MWCO, MilliporeSigma, Burlington, MA) to remove the free protein. Human CXCL12 ELISA kit (R&D systems) was used according to the manufacturer’s protocol to quantify total protein in the liposomes before centrifugation and free protein in the receiving tube after the centrifugation. CXCL12 loading efficiency (LE) was calculated using the following equation:

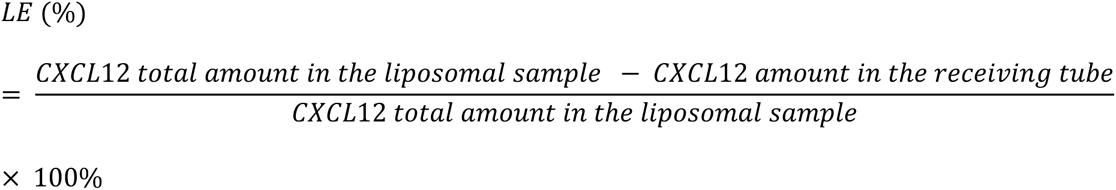

Lipofectamine 2000 was studied as blank liposomes. CXCL12-loaded liposomes were stored at 4°C until further use.

To form HA/Col II hydrogel, 5 mg/ml of HA solution in PBS (10 mM, pH 7.4) was mixed with 5 mg/ml of collagen II solution in 200 mM phosphate buffer (pH 7.4) (3/1, collagen II/HA, v/v). Following this, 200 mg/ml of 8-arm PEG succinimidyl glutarate (8-arm PEG) solution in PBS (10 mM, pH 7.4) was added into the mixture (1/13.5, v/v) and a HA/Col II hydrogel was formed following incubation at 37°C To form liposome/hydrogel composite (CXCL12-loaded or blank liposome-HA/Col II hydrogel), liposomes were accurately added into the phosphate buffer (200 mM, pH 7.4) to obtain the collagen II solution prior to mixing with other gel components (i.e., HA and PEG). The liposome/hydrogel composite was then formed when incubating the mixture at 37°C.

Gelation time of the liposome-HA/Col II hydrogel was assessed using a tube inversion method. Rheological properties of liposome-HA/Col II hydrogel were determined using a TA Discovery HR-2 rheometer (TA Instruments, New Castle, DE). The gel component mixture (800 μl) was immediately loaded on a sample plate that was pre-equilibrated at 25°C, and the cone shape geometry (40 mm, 2°) was lowered to a gap of 48 µm. Storage modulus of the liposome-HA/Col II hydrogel was determined via unconfined compression measurements with oscillation frequency ranging from 0.1 to 10 Hz at 5% strain at 37°C.

To evaluate swelling behavior of the hydrogel, disc-shaped liposome-embedded HA/Col II hydrogel samples were formed in a 96-well plate in triplicate. The sample was transferred into an Eppendorf tube and weighed on a microscale (Mettler Toledo, Greifensee, Switzerland) to obtain the initial weight (W_i_). The sample was incubated in an artificial cerebrospinal fluid (aCSF, Tocris Bioscience) at 37°C to allow free swelling for 48 hours. The sample was then removed and excess aCSF was gently removed. The fully swollen samples (W_s_) were recorded. Hydrogel swelling was calculated based on mass change percentage (%) using the following equation:

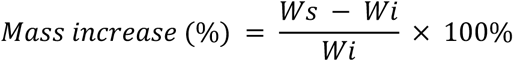

To determine CXCL-12 release in vitro, CXCL12-loaded liposome-HA/Col II hydrogel samples were pre-formed and incubated in 200 μl of aCSF. The samples were placed in a shaking water bath at 37°C with 45 rpm shaking speed. At predetermined time points, supernatants were collected and replenished with fresh aCSF. CXCL12 released amount was determined using a human CXCL12 ELISA kit (R&D systems) after breaking liposomes using 1% of Triton X-100. Cumulative CXCL-12 released amount was then calculated.

### In-vitro hydrogel gelation test

Gelation of the hydrogel was tested in an in-vitro environment that mimics the anatomy of the mouse brain. Matrigel (Corning, 354234) in DPBS was mixed and added to a 24-well plate, and this plate was incubated for at least 30 minutes to ensure the gelation of the Matrigel. Dextran, Oregon Green™ 488; 10 kDa, Anionic (D7170, Thermo Fisher Scientific) 25 mg/ml (1 ul) was mixed with 4 ul of DPBS. The diluted dextran (0.5 ul) was mixed with 100 ul of hydrogel components. The dextran-contained hydrogel component (2 ul) was injected into the pre-warmed Matrigel at -1.0 mm depth DV using a stereotactic injector with a 30G needle and Hamilton syringe at 0.5 ul/min. Right after the completion of the injection, the plate was imaged using Zeiss Apotome 2 with EGFP and a bright field at 10x magnification. This procedure was done with a tile mode (with tile size 10,000 um x 10,000 um and the number of tiles 224) and imaged at different time points (e.g., 30, 60, and 180 minutes).

### Cytokine array analysis

Proteome Profiler Mouse Cytokine Array kit (R&D) was used to detect the potential immune-inflammatory proteins using capture antibodies in an ELISA format, which are on a membrane with their predetermined locations. The brain lysate was prepared by homogenizing the right hemisphere of the mouse brain with the frontal lobe removed in 500 ul of pre-chilled DPBS with a protease inhibitor cocktail. Triton-X 100 (1%) was added to the homogenized cell lysate and stored at -80°C overnight. The thawed lysate was centrifuged at 10,000 xg for 5 minutes. Cytokine array assay was then performed on the supernatant of the mouse brain lysate by following the manufacturer’s instructions. The spots in the image of the array were quantified using an image software, FIJI, and the signal was normalized by the reference spots.

### In-vivo glioblastoma stem cell injection

All animal experiments were approved by Rhode Island Hospital’s Institutional Animal Care and Use Committee (IACUC) and conformed to the relevant regulatory standards and overseen by the institutional review board. Female and male NU/J homozygous mice (9 weeks old) were obtained from Jackson Laboratory (RRID:IMSR_JAX:002019) and used for the stereotactic injections. The animals were housed together until the day before the surgery and then housed individually. Animals receiving treatment were randomly assigned. ZsGreen expressing GSCs suspension 1x10^5^/ul in HPBS was prepared and loaded into a Hamilton syringe with a 30G needle. The tip of the needle was inserted into a 3.5 mm depth at a speed of 1.00 mm per minute. The needle was pulled back to 3.0 mm depth, and GSC suspension (2.0 ul) was injected at 0.5 ul/minute and rested for 15 minutes. The needle was pulled back at 1.00 mm per minute, and the hole was closed by Bonewax (Ethicon) and Vetbond (3M).

### In-vivo hydrogel implant

The experimental group animals received 10 µl of CXCL12-containing liposome (CL)-embedded hydrogel, while the control group animals received 10 µl of blank liposome (BL)-embedded hydrogel. For the CL animals, 6.9 µl of 5 mg/ml CXCL12-contained liposomes embedded collagen solution and 2.41 µL of 5 mg/ml HA in PBS solution were combined into a clean, sterile, and pre-chilled 1.5 ml Eppendorf tube. Following gentle vortex, 0.69 µL of 200 mg/mL 8-arm PEG was added to the tube. The hydrogel components were mixed gently and immersed in ice and subsequently injected into the mouse brain of the B6 mouse at coordinates of −2.0 mm AP and +1.5 mm ML relative to Bregma. Particularly, the syringe needle was lowered to -3.5 mm DV at a rate of 0.5 mm per minute, pulled up to -2.0 mm DV, and then injected with the hydrogel at 0.66 ul per minute. For the BL mice, the same procedure was followed except that 6.9 µl of 5 mg/ml liposome-contained collagen was used.

### Tissue clearing

The freshly harvested brains were fixed in 4% PFA at 4°C overnight. Brains were rinsed with pre-chilled PBS several times and transferred to pre-chilled PBS. Brains were cleared using X-CLARITYTM Hydrogel Solution Kit (Ledogos), according to the manufactured protocol, and stored in PBS at room temperature. The stock solution was prepared by mixing 2.5 g of X-CLARITY Polymerization Initiator (Cat# 13104, Logos Biosciences) in 10 ml 1X PBS to make a 25% (W/V) solution. Aliquots of 0.5 ml per brain were used and can be stored at -20°C for up to 6 months. After thawing the aliquot at 4°C or on ice, we diluted the solution 100 times in X-CLARITY Hydrogel Solution (Cat# 13103, Logos Biosciences). The brains were incubated in the hydrogel mixture at 4°C for 24 hours. The sample should be fully submerged in the hydrogel mixture.

### Hydrogel polymerization for 3D light sheet

The polymerization system’s settings were: Vacuum -90 kPa, Temperature 37°C, Timer 3 hours, and vessel type tube. The tube’s cap should not be tight to allow gas exchange during the procedure. After the incubation, the tubes were shaken for 1 minute to dissociate the hydrogel from the tube. The sample was then placed in a fresh tube and rinsed with 1X PBS multiple times to remove the excess hydrogel.

### Electrophoretic tissue clearing

The electrophoretic chamber was filled with Electrophoretic Tissue Clearing Solution (Cat# C13001, Logos Biosciences) before placing the sample holder in the chamber. The following settings were used for 16 to 18 hours to clear a whole brain: Current 1A, Temperature 37°C, and Pump speed 30 rpm. The samples were checked every 6 to 8 hours for clearing results. After removing the sample from the chamber, it was rinsed with 1X PBS several times. The sample appeared opaque and white in PBS.

### Light sheet microscope

The mouse brain was placed in 60% 2,2’-thiodiethanol solution overnight prior to imaging. The cleared brain was imaged using the LaVision Ultramicroscope II light-sheet microscope (Olympus). To image the entire brain, the image of the brain was acquired in a mosaic mode with 10% overlap and Z-stack mode with a light-sheet thickness of 4.05 um at magnification 2x. This image was analyzed by using the imaging software Imaris (Oxford Instruments). The tumor was defined based on the signals around the tumor injection site, particularly the tumor surface as contrast with the loss of signals in the surrounding area. The concentrated signals along the injection site were considered an artifact. The migrating cells were defined as dots with a local maximum outside the defined tumor and artifact. The migrating cells on top of the tumor were defined as the migrating cells above the middle of the tumor in the Z-axis, and the migrating cells on the bottom as those below that. For the mosaic 3D video, the images were taken with 10% overlap, and Imaris Stitcher was used to stitch 3D tile data and to create a video.

### Visualization of hydrogel distribution in the brain

The HA/Col II hydrogel components was prepared using the same formulation as described above. Dextran (25 mg/ml, 1.1 μl) was added to the mixture to form the dextran-loaded hydrogel, followed by the addition of a 200 mg/ml 8-arm PEG solution in PBS (10 mM, pH 7.4) (1/13.5, v/v). The dextran-loaded hydrogel (2 μl) was then injected into the mouse brain using a stereotactic device, at a speed of 0.666 μl/minute to a depth of 2 mm. One hour after the injection, the mouse was sacrificed and the brain was harvested, fixed in 4% PFA overnight, and transferred to pre-chilled PBS within 24 hours. The brain went through the tissue clearing, hydrogel polymerization, and electrophoretic tissue clearing steps. Subsequently, the cleared brain was washed with PBST and transferred to 1:300 Anti-Actin, α-Smooth Muscle - Cy3™ antibody (Millipore Sigma) in PBST for staining of the blood vessels. The sample was washed with distilled water to remove phosphate. The sample was incubated at X-CLARITY™ Mounting Solution for 1 hour at room temperature, followed by a rinse with fresh X-CLARITY™ Mounting Solution and additional incubation for 1-2 hours.

## List of Supplementary Materials

**Supplementary Figure 1**

**Supplementary Figure 2**

## Supporting information

Supplemental figure legens

## Acknowledgments

We would like to thank Drs. Steven Toms and Atom Sarkar for providing human glioblastoma tumor samples in support of our research. We would also like to thank Drs. Margot Martinez Moreno and Emily Arner for performing proof-of-concept experiments during the initial conceptualization phase of this project. We would like to thank Rajeev Kant and Amanda Khoo for their help in designing the PDMS device. We would like to thank the Neurobiology Imaging Facility at Harvard Medical School for consultation and instrument availability that supported this work. This facility is supported in part by the Neural Imaging Center as part of an NINDS P30 Core Center grant #NS072030. We thank the Lentivirus Construct Core of the Center for Stem Cells and Aging at Brown University supported by a COBRE grant from the National Institutes of Health (P20 GM119943).

## Funding

Department of Neurosurgery, Brown University

Private donations to the Tapinos Lab

Warren Alpert Foundation

## Author contributions

Conceptualization: NT

Methodology: NT, JS, YS

Investigation: YS, DM-T, SM, WY, LX, BA, NP, EF, AF

Visualization: YS, NT

Funding acquisition: NT, JS

Supervision: NT, JS

Writing – original draft: YS, NT

NT Writing – review & editing: NT, JS

## Data and materials availability

The RNA-seq data discussed in this manuscript have been deposited in NCBI’s Gene Expression Omnibus and are accessible through GEO Series accession number GSE219288.

## References and Notes

1. R. Stupp, W. P. Mason, M. J. van den Bent, M. Weller, B. Fisher, M. J. Taphoorn, K. Belanger, A. A. Brandes, C. Marosi, U. Bogdahn, J. Curschmann, R. C. Janzer, S. K. Ludwin, T. Gorlia, A. Allgeier, D. Lacombe, J. G. Cairncross, E. Eisenhauer, R. O. Mirimanoff, R. European Organisation for, T. Treatment of Cancer Brain, G. Radiotherapy, G. National Cancer Institute of Canada Clinical Trials, Radiotherapy plus concomitant and adjuvant temozolomide for glioblastoma. N Engl J Med 352, 987–996 (2005).

2. R. Stupp, D. C. Weber, The role of radio- and chemotherapy in glioblastoma. Onkologie 28, 315–317 (2005).

3. S. K. Singh, I. D. Clarke, M. Terasaki, V. E. Bonn, C. Hawkins, J. Squire, P. B. Dirks, Identification of a cancer stem cell in human brain tumors. Cancer Res 63, 5821–5828 (2003).

4. S. K. Singh, C. Hawkins, I. D. Clarke, J. A. Squire, J. Bayani, T. Hide, R. M. Henkelman, M. D. Cusimano, P. B. Dirks, Identification of human brain tumour initiating cells. Nature 432, 396–401 (2004).

5. J. Lee, S. Kotliarova, Y. Kotliarov, A. Li, Q. Su, N. M. Donin, S. Pastorino, B. W. Purow, N. Christopher, W. Zhang, J. K. Park, H. A. Fine, Tumor stem cells derived from glioblastomas cultured in bFGF and EGF more closely mirror the phenotype and genotype of primary tumors than do serum-cultured cell lines. Cancer Cell 9, 391–403 (2006).

6. P. Soni, S. Qayoom, N. Husain, P. Kumar, A. Chandra, B. K. Ojha, R. K. Gupta, CD24 and Nanog expression in Stem Cells in Glioblastoma: Correlation with Response to Chemoradiation and Overall Survival. Asian Pac J Cancer Prev 18, 2215–2219 (2017).

7. E. C. Collin, S. Grad, D. I. Zeugolis, C. S. Vinatier, J. R. Clouet, J. J. Guicheux, P. Weiss, M. Alini, A. S. Pandit, An injectable vehicle for nucleus pulposus cell-based therapy. Biomaterials 32, 2862–2870 (2011).

8. L. Xie, W. Yue, K. Ibrahim, J. Shen, A Long-Acting Curcumin Nanoparticle/In Situ Hydrogel Composite for the Treatment of Uveal Melanoma. Pharmaceutics 13, (2021).

9. L. J. Nesti, W. J. Li, R. M. Shanti, Y. J. Jiang, W. Jackson, B. A. Freedman, T. R. Kuklo, J. R. Giuliani, R. S. Tuan, Intervertebral disc tissue engineering using a novel hyaluronic acid-nanofibrous scaffold (HANFS) amalgam. Tissue Eng Part A 14, 1527–1537 (2008).

10. Y. R. Zou, A. H. Kottmann, M. Kuroda, I. Taniuchi, D. R. Littman, Function of the chemokine receptor CXCR4 in haematopoiesis and in cerebellar development. Nature 393, 595–599 (1998).

11. J. B. Rubin, A. L. Kung, R. S. Klein, J. A. Chan, Y. Sun, K. Schmidt, M. W. Kieran, A. D. Luster, R. A. Segal, A small-molecule antagonist of CXCR4 inhibits intracranial growth of primary brain tumors. Proc Natl Acad Sci U S A 100, 13513–13518 (2003).

12. N. Goffart, J. Kroonen, E. Di Valentin, M. Dedobbeleer, A. Denne, P. Martinive, B. Rogister, Adult mouse subventricular zones stimulate glioblastoma stem cells specific invasion through CXCL12/CXCR4 signaling. Neuro Oncol 17, 81–94 (2015).

13. A. Bajetto, F. Barbieri, A. Dorcaratto, S. Barbero, A. Daga, C. Porcile, J. L. Ravetti, G. Zona, R. Spaziante, G. Corte, G. Schettini, T. Florio, Expression of CXC chemokine receptors 1-5 and their ligands in human glioma tissues: role of CXCR4 and SDF1 in glioma cell proliferation and migration. Neurochem Int 49, 423–432 (2006).

14. J. C. Gutjahr, K. S. Crawford, D. R. Jensen, P. Naik, F. C. Peterson, G. P. B. Samson, D. F. Legler, J. Duchene, C. T. Veldkamp, A. Rot, B. F. Volkman, The dimeric form of CXCL12 binds to atypical chemokine receptor 1. Sci Signal 14, (2021).

15. Y. Shi, D. J. Riese, 2nd, J. Shen, The Role of the CXCL12/CXCR4/CXCR7 Chemokine Axis in Cancer. Front Pharmacol 11, 574667 (2020).

16. A. R. Pombo Antunes, I. Scheyltjens, F. Lodi, J. Messiaen, A. Antoranz, J. Duerinck, D. Kancheva, L. Martens, K. De Vlaminck, H. Van Hove, S. S. Kjolner Hansen, F. M. Bosisio, K. Van der Borght, S. De Vleeschouwer, R. Sciot, L. Bouwens, M. Verfaillie, N. Vandamme, R. E. Vandenbroucke, O. De Wever, Y. Saeys, M. Guilliams, C. Gysemans, B. Neyns, F. De Smet, D. Lambrechts, J. A. Van Ginderachter, K. Movahedi, Single-cell profiling of myeloid cells in glioblastoma across species and disease stage reveals macrophage competition and specialization. Nat Neurosci 24, 595–610 (2021).

17. M. J. Rowland, C. C. Parkins, J. H. McAbee, A. K. Kolb, R. Hein, X. J. Loh, C. Watts, O. A. Scherman, An adherent tissue-inspired hydrogel delivery vehicle utilised in primary human glioma models. Biomaterials 179, 199–208 (2018).

18. I. Dotan, L. Werner, S. Vigodman, S. Weiss, E. Brazowski, N. Maharshak, O. Chen, H. Tulchinsky, Z. Halpern, H. Guzner-Gur, CXCL12 is a constitutive and inflammatory chemokine in the intestinal immune system. Inflamm Bowel Dis 16, 583–592 (2010).

19. E. M. Garcia-Cuesta, C. A. Santiago, J. Vallejo-Diaz, Y. Juarranz, J. M. Rodriguez-Frade, M. Mellado, The Role of the CXCL12/CXCR4/ACKR3 Axis in Autoimmune Diseases. Front Endocrinol (Lausanne) 10, 585 (2019).

20. E. E. McCandless, Q. Wang, B. M. Woerner, J. M. Harper, R. S. Klein, CXCL12 limits inflammation by localizing mononuclear infiltrates to the perivascular space during experimental autoimmune encephalomyelitis. J Immunol 177, 8053–8064 (2006).

21. E. Lee, J. Choi, Y. Jo, J. Y. Kim, Y. J. Jang, H. M. Lee, S. Y. Kim, H. J. Lee, K. Cho, N. Jung, E. M. Hur, S. J. Jeong, C. Moon, Y. Choe, I. J. Rhyu, H. Kim, W. Sun, ACT-PRESTO: Rapid and consistent tissue clearing and labeling method for 3-dimensional (3D) imaging. Sci Rep 6, 18631 (2016).

22. K. Chung, J. Wallace, S. Y. Kim, S. Kalyanasundaram, A. S. Andalman, T. J. Davidson, J. J. Mirzabekov, K. A. Zalocusky, J. Mattis, A. K. Denisin, S. Pak, H. Bernstein, C. Ramakrishnan, L. Grosenick, V. Gradinaru, K. Deisseroth, Structural and molecular interrogation of intact biological systems. Nature 497, 332–337 (2013).

23. A. A. Mughal, L. Zhang, A. Fayzullin, A. Server, Y. Li, Y. Wu, R. Glass, T. Meling, I. A. Langmoen, T. B. Leergaard, E. O. Vik-Mo, Patterns of Invasive Growth in Malignant Gliomas-The Hippocampus Emerges as an Invasion-Spared Brain Region. Neoplasia 20, 643–656 (2018).

24. J. P. Bruggeman, B. J. de Bruin, C. J. Bettinger, R. Langer, Biodegradable poly(polyol sebacate) polymers. Biomaterials 29, 4726–4735 (2008).

25. H. J. Scherer, A Critical Review: The Pathology of Cerebral Gliomas. J Neurol Psychiatry 3, 147–177 (1940).

26. M. Ehtesham, J. A. Winston, P. Kabos, R. C. Thompson, CXCR4 expression mediates glioma cell invasiveness. Oncogene 25, 2801–2806 (2006).

27. D. Zagzag, M. Esencay, O. Mendez, H. Yee, I. Smirnova, Y. Huang, L. Chiriboga, E. Lukyanov, M. Liu, E. W. Newcomb, Hypoxia- and vascular endothelial growth factor-induced stromal cell-derived factor-1alpha/CXCR4 expression in glioblastomas: one plausible explanation of Scherer’s structures. Am J Pathol 173, 545–560 (2008).

28. J. M. Munson, R. V. Bellamkonda, M. A. Swartz, Interstitial flow in a 3D microenvironment increases glioma invasion by a CXCR4-dependent mechanism. Cancer Res 73, 1536–1546 (2013).

29. A. Bajetto, F. Barbieri, A. Pattarozzi, A. Dorcaratto, C. Porcile, J. L. Ravetti, G. Zona, R. Spaziante, G. Schettini, T. Florio, CXCR4 and SDF1 expression in human meningiomas: a proliferative role in tumoral meningothelial cells in vitro. Neuro Oncol 9, 3–11 (2007).

30. B. M. Woerner, N. M. Warrington, A. L. Kung, A. Perry, J. B. Rubin, Widespread CXCR4 activation in astrocytomas revealed by phospho-CXCR4-specific antibodies. Cancer Res 65, 11392–11399 (2005).

31. M. Sciaccaluga, G. D’Alessandro, F. Pagani, G. Ferrara, N. Lopez, T. Warr, P. Gorello, A. Porzia, F. Mainiero, A. Santoro, V. Esposito, G. Cantore, E. Castigli, C. Limatola, Functional cross talk between CXCR4 and PDGFR on glioblastoma cells is essential for migration. PLoS One 8, e73426 (2013).

32. R. Wurth, A. Bajetto, J. K. Harrison, F. Barbieri, T. Florio, CXCL12 modulation of CXCR4 and CXCR7 activity in human glioblastoma stem-like cells and regulation of the tumor microenvironment. Front Cell Neurosci 8, 144 (2014).

33. N. Goffart, A. Lombard, F. Lallemand, J. Kroonen, J. Nassen, E. Di Valentin, S. Berendsen, M. Dedobbeleer, E. Willems, P. Robe, V. Bours, D. Martin, P. Martinive, P. Maquet, B. Rogister, CXCL12 mediates glioblastoma resistance to radiotherapy in the subventricular zone. Neuro Oncol 19, 66–77 (2017).

34. J. P. Zepecki, K. M. Snyder, M. M. Moreno, E. Fajardo, A. Fiser, J. Ness, A. Sarkar, S. A. Toms, N. Tapinos, Regulation of human glioma cell migration, tumor growth, and stemness gene expression using a Lck targeted inhibitor. Oncogene, (2018).

35. C. A. Schneider, W. S. Rasband, K. W. Eliceiri, NIH Image to ImageJ: 25 years of image analysis. Nat Methods 9, 671–675 (2012).

36. D. Kim, J. M. Paggi, C. Park, C. Bennett, S. L. Salzberg, Graph-based genome alignment and genotyping with HISAT2 and HISAT-genotype. Nat Biotechnol 37, 907–915 (2019).

37. M. I. Love, W. Huber, S. Anders, Moderated estimation of fold change and dispersion for RNA-seq data with DESeq2. Genome Biol 15, 550 (2014).

38. A. Subramanian, P. Tamayo, V. K. Mootha, S. Mukherjee, B. L. Ebert, M. A. Gillette, A. Paulovich, S. L. Pomeroy, T. R. Golub, E. S. Lander, J. P. Mesirov, Gene set enrichment analysis: a knowledge-based approach for interpreting genome-wide expression profiles. Proc Natl Acad Sci U S A 102, 15545–15550 (2005).

39. P. Langfelder, S. Horvath, WGCNA: an R package for weighted correlation network analysis. BMC Bioinformatics 9, 559 (2008).

40. J. A. Gustavsen, S. Pai, R. Isserlin, B. Demchak, A. R. Pico, RCy3: Network biology using Cytoscape from within R. F1000Res 8, 1774 (2019).

