## Supplemental figure legens for "Engineering GliaTrap: a biodegradable non-swelling hydrogel with tuned release of CXCL12 to attract migrating glioblastoma cells"

**Supplementary Figure 1:**

A) MTS assay of CXCl12-treated GSCs shows that CXCL12 does not induce significant increase in proliferation (n=3, Student's t-test). B) Volcano plot of differential gene expression analysis on CXCL12-treated GSCs. Genes with |fold change|>1.5 and p-value<0.05 were considered as significant (Red dots: Up-regulated genes, blue dots: down-regulated genes, grey: genes with no-significant difference). C) Geneset Enrichment Analysis (GSEA) showed that the Chemotaxis & Epithelial Mesenchymal Transition (EMT) gene signatures are enriched. The right panel shows the genes that are up-regulated in CXCL12-treated GSCs. D) GSEA showed that the GBM Proneural signature and Mesenchymal signature are enriched. E) Plot of WGCNA on CXCL12 "high" GSCs using Cytoscape shows that CXCL12 connects with CD248, SRGN, and ABI3BP.

**Supplementary Figure 2:**

A) Setting of Dextran-loaded hydrogel injection into pre-warmed Matrigel using the stereotactic device (green circle shows the Hamilton syringe that injects hydrogel into pre-warmed Matrigel) B) Images were taken 30 minutes after injecting the Dextran + hydrogel group into pre-warmed Matrigel to observe the release of Dextran from the hydrogel. The images indicate that Dextran was released slower from the hydrogel compared to Dextran alone. C) Images of the mouse brain injected with each hydrogel (each circle shows the hydrogel injection site).
