## Supplementary figures and images for "Engineering GliaTrap: a biodegradable non-swelling hydrogel with tuned release of CXCL12 to attract migrating glioblastoma cells"

### Supplementary Figures

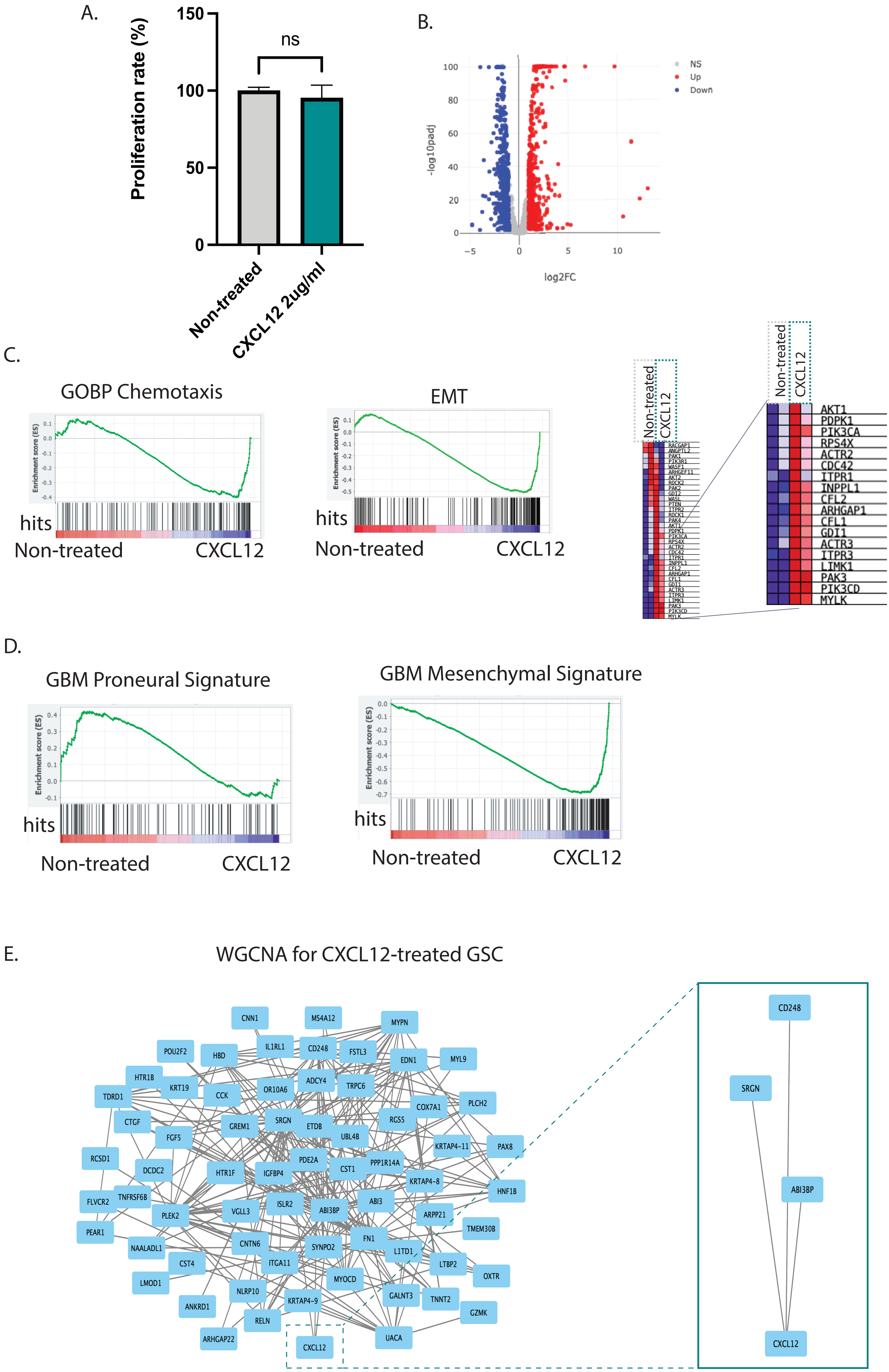

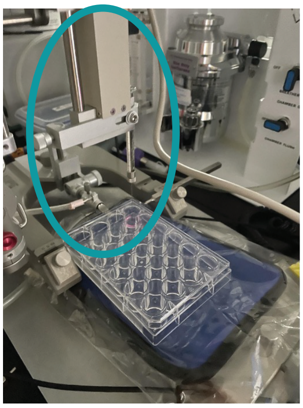

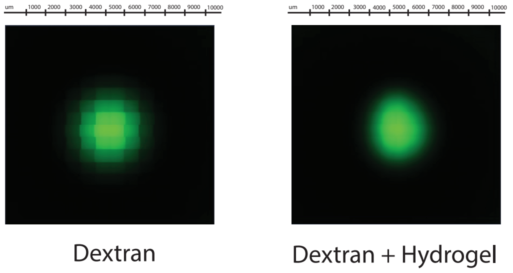

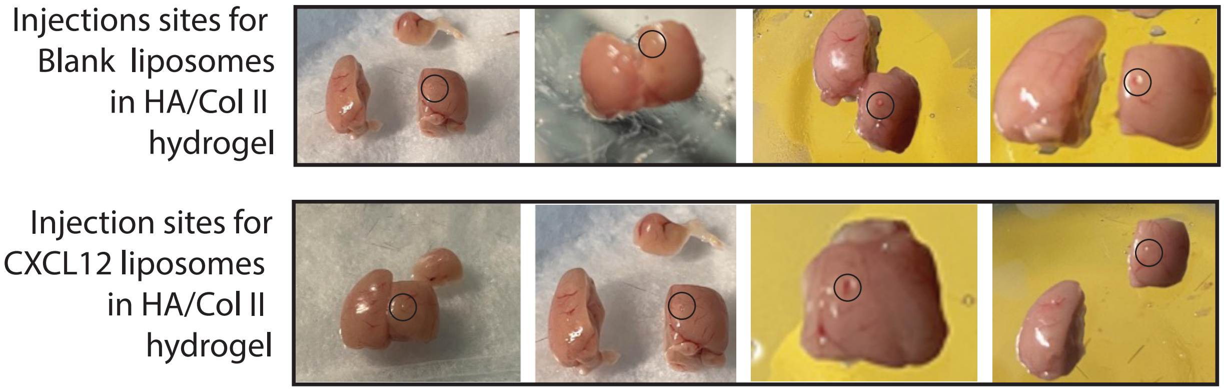
